# KMT2D-NOTCH Mediates Coronary Abnormalities in Hypoplastic Left Heart Syndrome

**DOI:** 10.1101/2021.08.30.457716

**Authors:** Zhiyun Yu, Xin Zhou, Ziyi Liu, Victor Pastrana-Gomez, Yu Liu, Minzhe Guo, Lei Tian, Timothy J. Nelson, Nian Wang, Seema Mital, David Chitayat, Joseph C. Wu, Marlene Rabinovitch, Sean M. Wu, Michael P. Snyder, Yifei Miao, Mingxia Gu

## Abstract

Hypoplastic left heart syndrome (HLHS) is a severe form of single ventricle congenital heart disease characterized by the underdevelopment of the left ventricle. Early serial postmortem examinations revealed high rate of coronary artery abnormalities in HLHS fetal hearts, such as thickened wall, kinking arteries and ventriculo-coronary arterial connection. However, it is unclear if there is an intrinsic defect in the HLHS coronary vessels and what the underlying molecular mechanism is.

Here, we profiled both human fetal heart with an underdeveloped left ventricle (ULV) and ECs differentiated from induced pluripotent stem cells (iPSCs) derived from HLHS patients at single cell resolution. CD144^+^/NPR3^-^vascular ECs were selected and further classified as venous, arterial and late arterial subclusters. To study the arterial EC phenotype, we specifically generated iPSC-arterial ECs (AECs, CD34^+^CDH5^+^CXCR4^+^NT5E^-/low^) derived from three HLHS patients and three age-matched healthy controls.

Gene ontology analysis revealed that ULV late arterial EC subcluster showed specific defects in endothelial development, proliferation, and Notch signaling compared to control. Consistently, HLHS iPSCs exhibited impaired AEC differentiation shown as the reduced CXCR4^+^NT5E^-/low^ AEC progenitor population. Mature HLHS iPSC-AECs also exhibited increased G0/G1 cell cycle arrest with decreased expression of cell cycle related genes (e.g., Ki67, CCND1/2). Additionally, NOTCH targeted genes (e.g., DLL4, HEY1, GJA5) were found suppressed in both ULV AECs and HLHS iPSC-AECs compared to control. We also found the HLHS de novo mutation gene KMT2D directly regulated the transcription of NOTCH targeted genes participating in arterial differentiation and cell proliferation, contributing to the HLHS AEC dysfunctionalities. Intriguingly, the treatment of NOTCH ligand JAG1 improved cell proliferation of HLHS AECs and upregulated G1/S transition genes downstream of NOTCH pathway.

In summary, our results revealed that KMT2D directly regulated transcription activity of NOTCH signaling, contributing to the poor differentiation and low proliferation of HLHS coronary AECs.

## Main text

Hypoplastic left heart syndrome (HLHS) is a severe form of single ventricle congenital heart disease (CHD) characterized by the underdevelopment of the left ventricle, mitral valve, aortic valve, and ascending aorta. Coronary arterial abnormalities such as thickened wall, kinking arteries, and coronary arterial fistulous communications have been revealed by postmortem examinations^1^, which may impact ventricular development and intra-cardiac hemodynamics, leading to a poor prognosis after surgical palliation^1^. However, the intrinsic defect in coronary vessels and its genetic basis remain unclear. Through single-cell RNA sequencing (scRNA-seq) analysis of human fetal heart with an underdeveloped left ventricle (ULV) and ECs differentiated from induced pluripotent stem cells (iPSCs) with HLHS, we uncovered an abnormal population of coronary arterial ECs with loss of arterial features and decreased proliferation, which were attributed to HLHS *de novo* mutation (DNM) *KMT2D* mediated NOTCH signaling defect.

To reveal the transcriptomic changes in HLHS coronary vessels, heart ECs (CDH5^+^) were enriched from dissociated human fetal heart (healthy control vs. ULV) (**A**) and iPSC-derived ECs (iPSC-ECs, healthy control vs. HLHS)^2^ (**Online A**) and subjected to scRNA-seq. We first excluded the endocardial population (NPR3^+^) by selecting CDH5^+^/NPR3^-^ vascular ECs^2^ for downstream analysis. Out of the six sub-clusters in fetal heart vascular ECs, we focused on four clusters (Cluster 0-3) containing predominate vascular ECs and excluded those in transitional states. EC subtypes were annotated by multiple cell-type-specific markers, such as vein (*NR2F2*^+^, Cluster 0&2), artery (*MECOM*^+^, Cluster 1&3) (**Online A**), and late artery (*GJA5*^+^, Cluster 1)^3^ (**A**). iPSC-vascular ECs were heterogeneous and classified into four sub-clusters. The majority of the cells (Cluster 0&1) showed arterial characteristics (high *MECOM, GJA5*, and low *NR2F2*), possibly due to the high VEGF in the differentiation medium that induced an arterial cell fate^2^ (**Online A**). Gene ontology (GO) analysis was performed based on differentially expressed genes (DEGs) between control and ULV vascular ECs (**A**). Compared with control, ULV showed defects in general EC functions such as cell junction organization and EC migration. Notably, C1 late arterial cluster exhibited cell-type specific defects in endothelium development, EC proliferation, artery morphogenesis, and Notch signaling, which were not observed in other EC subclusters. These results intrigued us to further focus on understanding the coronary arterial ECs (AECs) abnormalities in HLHS.

Next, we generated pure AECs from three HLHS and age-matched control iPSC lines^2^ using a published protocol^4^. HLHS iPSCs exhibited impaired AEC differentiation as evidenced by the reduced CXCR4^+^NT5E^-/low^ AEC progenitors compared to the control (**Ba**). Mature AECs were further enriched by CXCR4^+^ cell sorting. HLHS AECs demonstrated impaired proliferation with increased G0/G1 and decreased S/G2/M cell percentage (**Ba**), accompanied by the downregulation of *Ki67* and G1/S transition genes *(CCND1/2),* and the upregulation of G1/S transition inhibitor *CDKN2A/P16* (**Online B**). This was further validated in HLHS human fetal heart tissue showing reduced proliferative AECs labeled by Ki67 (**Bb**). Consistent with GO analysis showing NOTCH defect in ULV AECs (**A**), several critical NOTCH targeted arterial genes (*GJA5*, *DLL4, HEY1)* were suppressed in AECs from ULV (Cluster 1) and HLHS iPSC-ECs (Cluster 0&1) (**Online C**). Decreased expression of NOTCH targeted genes and NOTCH intracellular domain (NICD) were also observed in HLHS iPSC-AECs (**Ca**) and fetal heart tissue (**Cb**).

Previously, we revealed that the majority of the HLHS DNMs encoded genes were highly enriched in endocardial and endothelial populations in human fetal heart^2^. To further understand the genetic underpinnings in the HLHS cases we studied, we first examined the expression levels of several key chromatin remodelers harboring HLHS DNMs. We identified five genes (e.g., *FOXM1, KMT2D)* that significantly reduced in HLHS iPSC-ECs (**Da**). Among them, KMT2D, a lysine methyltransferase, favors NICD-RBPJ complex mediated gene activation by maintaining a permissive chromatin status via catalyzing H3K4me1, me2 and me3^5^. Interestingly, knockdown of *KMT2D* in primary coronary ECs suppressed the expression of NOTCH related genes (**Online D**). KMT2D protein level was also reduced in HLHS coronary arteries (**Db**). Additionally, ChIP-qPCR revealed reduced KMT2D binding capacity and H3K4me2 signals to the promoter loci of several NOTCH targeted genes in mature HLHS AECs (**E**), which are critical in maintaining arterial characteristics *(DLL4, HEY1/2, HES1)* and cell proliferation *(CCND1).* Intriguingly, the treatment of NOTCH ligand Jag1 and Dll1 improved cell proliferation of HLHS AECs and upregulated G1/S transition genes downstream of the NOTCH pathway (**F**).

In summary, our study revealed that KMT2D-mediated NOTCH defect contributed to the coronary AECs abnormalities in HLHS. The NOTCH-related defects were more pronounced in coronary AECs compared with other cardiac cell types. Disruption of NOTCH signaling led to impaired proliferation and maintenance of arterial features in HLHS AECs, which may partially explain the decreased vascular density and coronary artery malformation in HLHS fetal heart^1^. Notably, NOTCH ligands rescued HLHS associated gene expression abnormalities and cellular phenotypes, providing a potential therapeutic target.

## Footnotes

Consent for iPSC generation was obtained from both control and patients under approved IRBs: Mayo Clinic: 10-006845; Stanford: IRB 5443. Tissue collection and use in the research were approved by the University of Washington: IRB STUDY00000380. Human tissue sections were obtained under approved IRBs: Hospital for Sick Children IRB 1000011284; Mount Sinai Hospital REB# 08-0009-E.

## Data sharing

Raw data and complete methods can be made available upon request from the corresponding author. Single-cell RNA sequencing Seurat object have been deposited in the GEO database under accession number GSE138979. Online supplemented figure can be found in Biorxiv version.

## The Correspondence

Yifei Miao, PhD, Cincinnati Children’s Hospital Medical Center, Room R4.464; MLC 7029, 3333 Burnet Ave, Cincinnati, OH 45229

Mingxia Gu, MD, PhD, Cincinnati Children’s Hospital Medical Center, Room R4.467; MLC 7029, 3333 Burnet Ave, Cincinnati, OH 45229

## Affiliations

Department of Genetics (X.Z., M.S.), Cardiovascular Institute (X.Z., Y.L., L.T., S.M.W., J.C.W., M.R., Y.M., M.G.), Department of Pediatrics, Division of Pediatric Cardiology (S.M.W., M.R., Y.M., M.G.), Institute for Stem Cell Biology and Regenerative Medicine & Department of Medicine, Division of Cardiovascular Medicine (S.M.W., J.C.W.,), Department of Radiology (J.C.W.), Stanford University School of Medicine, Stanford, CA.

Division of Pediatric Cardiology, Department of Pediatric and Adolescent Medicine (T.J.N.), Department of Molecular Pharmacology & Experimental Therapeutics (T.J.N.), General Internal Medicine and Transplant Center, Department of Internal Medicine (T.J.N.), Center for Regenerative Medicine (T.J.N.), Mayo Clinic, Rochester, MN.

Department of Radiology and Imaging Sciences (N.W.), Stark Neurosciences Research Institute (N.W.), Indiana University, Indianapolis, IN.

Department of Pediatrics, Hospital for Sick Children (S.M., D.C.), The Prenatal Diagnosis and Medical Genetics Program, Department of Obstetrics and Gynecology, Mount Sinai Hospital (D.C.), University of Toronto, Toronto, Canada

Perinatal Institute, Division of Pulmonary Biology (Z.Y., Z.L., M.Z.G., Y.M., M.G.), Center for Stem Cell and Organoid Medicine, CuSTOM, Division of Developmental Biology (Z.Y., Z.L., Y.M., M.G.), Cincinnati Children’s Hospital Medical Center, Cincinnati, OH. University of Cincinnati School of Medicine (Z.Y., M.Z.G., Y.M., M.G.), Cincinnati, OH.

## Acknowledgments

We thank Drs. Kyle Loh, Nanhua Zhang for providing intellectual consultant to the project. We also thank ReGen Theranostics, Inc Rochester, MN as the manufacturer for the iPSC lines.

---

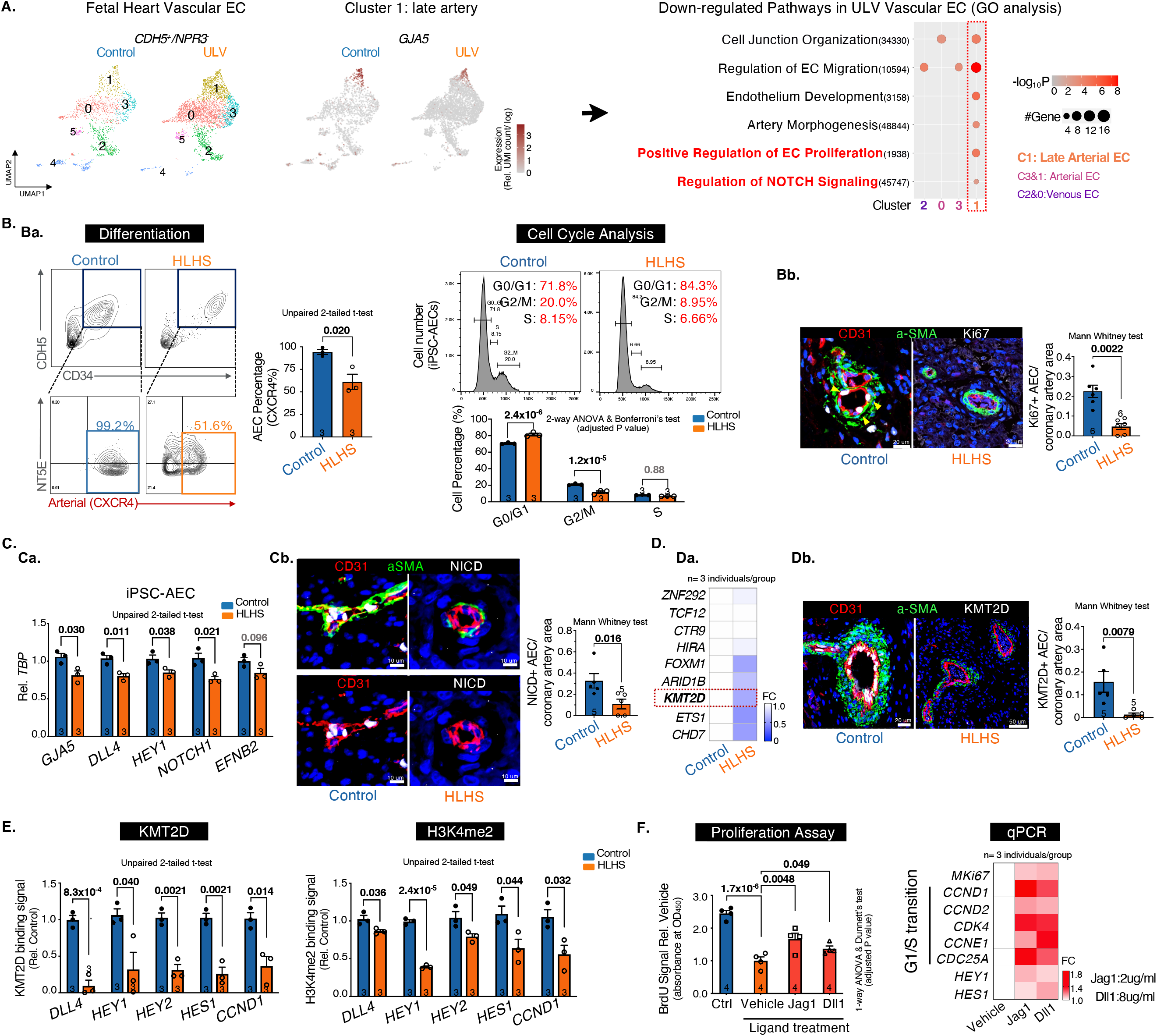
**A. scRNA-seq analysis of vascular ECs *(CDH5^+^INPR3^-^*).** UMAP projection of Control vs. ULV human fetal left heart vascular ECs were shown on the left. Cluster1: *GJA5^+^* late artery. Cluster4: venous-to-endocardial transitional population. N=1 per group. Then Gene Ontology analysis were performed on each cluster and showed downregulated pathways in ULV vascular ECs vs. control. GO terms: adjusted p-value < 0.05. UMI: Unique Molecular Identifier. **B. Artery differentiation and proliferation were impaired in HLHS. Ba**, flow cytometry analysis of iPSC-AEC progenitors percentage and their cell-cycle status. **Bb**, Immunostaining and quantification of Ki67^+^ arteries in human heart tissue. aSMA: smooth muscle actin. **C. NOTCH pathway was suppressed in HLHS AECs. Ca**, qPCR of NOTCH genes in iPSC-AECs. **Cb**, Immunostaining and quantification of NICD^+^ arteries in human heart tissue. **D. KMT2D expression was decreased in HLHS AECs. Da**, qPCR of HLHS DNM gene expression in iPSC-ECs. **Db**, Immunostaining and quantification of KMT2D^+^ arteries in human heart tissue. **E. KMT2D binding capacity and H3K4me2 signal were reduced in HLHS iPSC-AECs.** (Evaluated via ChIP-qPCR) **F. NOTCH ligands treatment improved the proliferation of HLHS iPSC-AECs.** Both BrdU proliferation assay and qPCR showed improved proliferation in HLHS iPSC-AECs. N=4 technical repeats. FC: fold change normalized to control. **Statistics** (GraphPad Prism 9.3.1): Based on additional literature support from similar studies, our samples fit normal distribution. Parametric test: unpaired 2-tailed t-test (2 groups), ANOVA (>2 groups) with post hoc tests as indicated; non-parametric test: Mann-Whitney (2 groups). Mean±SEM; n= biological replicates as indicated.

