## Supplementary material for "KMT2D-NOTCH Mediates Coronary Abnormalities in Hypoplastic Left Heart Syndrome": Online Figure

**Online Figures. Yu et al., Circulation Research, 2022.**

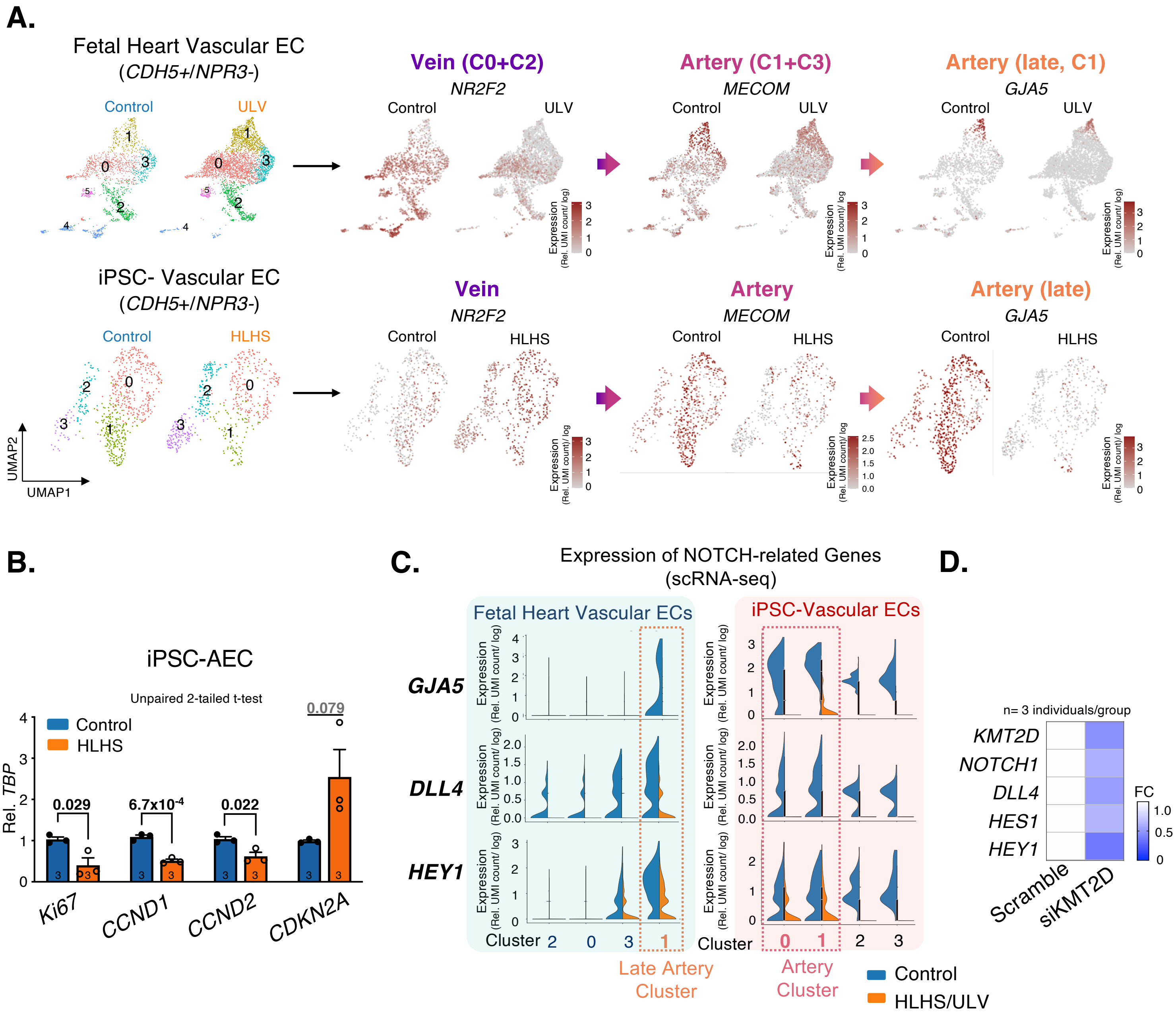


**Online Figure legend:**

**A. scRNA-seq analysis of vascular ECs (*CDH5*$^{+}$/*NPR3*$^{-}$). Top**: Control vs. ULV human fetal left heart vascular ECs. Cluster0+Cluster2: *NR2F2*$^{+}$ vein, Cluster1+Cluster3: *MECOM*$^{+}$ artery, Cluster1: *GJA5*$^{+}$ late artery. **Bottom**: Control vs. HLHS iPSC-vascular ECs. EC sub-clusters were labeled by the subtype markers. N=1 per group. UMI: Unique Molecular Identifier.

**B. Artery differentiation and proliferation were impaired in HLHS.** qPCR of cell cycle markers in control vs. HLHS iPSC-AECs.

**C. NOTCH pathway was suppressed in HLHS AECs.** Violin plots of NOTCH-arterial genes in vascular ECs from control vs. ULV fetal heart (left) and control vs. HLHS iPSC-ECs (right).

**D. KMT2D regulated the expression of NOTCH pathway genes in coronary ECs.** qPCR of NOTCH pathway genes in primary coronary ECs with *KMT2D* knockdown.

**Statistics** (GraphPad Prism 9.3.1): Based on additional literature support from similar studies, our samples fit normal distribution. Parametric test: unpaired 2-tailed t-test (2 groups). Mean±SEM; n= biological replicates as indicated.
